# Genetic Modifiers of Pathogenic LRRK2 G2019S Neurodegeneration in *Drosophila*

**DOI:** 10.1101/331991

**Authors:** Sierra Lavoy, Vinita G. Chittoor-Vinod, Clement Y. Chow, Ian Martin

## Abstract

Disease phenotypes can be highly variable among individuals with the same pathogenic mutation. There is increasing evidence that background genetic variation is a strong driver of disease variability in addition to the influence of environment. To understand the genotype-phenotype relationship that determines the expressivity of a pathogenic mutation, a large number of backgrounds must be studied. This can be efficiently achieved using model organism collections such as the *Drosophila* Genetic Reference Panel (DGRP). Here, we used the DGRP to assess the variability of locomotor dysfunction in a LRRK2 G2019S *Drosophila melanogaster* model of Parkinson’s disease. We find substantial variability in the LRRK2 G2019S locomotor phenotype in different DGRP backgrounds. A genome-wide association study for candidate genetic modifiers reveals 177 genes that drive wide phenotypic variation, including 19 top association genes. Genes involved in the outgrowth and regulation of neuronal projections are enriched in these candidate modifiers. RNAi functional testing of the top association and neuronal projection-related genes reveals that *pros, pbl, ct* and *CG33506* significantly modify age-related dopamine neuron loss and associated locomotor dysfunction in the *Drosophila* LRRK2 G2019S model. These results demonstrate how natural genetic variation can be used as a powerful tool to identify genes that modify disease-related phenotypes. We report novel candidate modifier genes for LRRK2 G2019S that may be used to interrogate the link between LRRK2, neurite regulation and neuronal degeneration in Parkinson’s disease.

## Introduction

Parkinson’s disease (PD) is an age-associated neurodegenerative disease of unknown cause. Established therapeutic approaches provide symptomatic relief, but do not modify the underlying disease (1). Recently, the contribution of genetic and epigenetic factors to PD development has been increasingly recognized (2). This follows the identification of pathogenic mutations in genes including *SNCA* (*Synuclein alpha*), *LRRK2 (leucine-rich repeat kinase 2), PINK1 (PTEN-induced putative kinase 1), parkin,* and *DJ-1,* along with numerous risk susceptibility loci found in idiopathic PD via genome-wide association studies (1, 2). LRRK2, which is linked to autosomal-dominant familial PD, has received much attention because (i) LRRK2-linked familial PD clinically and pathologically recapitulates late-onset idiopathic disease, (ii) the disease-causing G2019S mutation and multiple risk variants at the LRRK2 locus are prevalent in idiopathic PD, and (iii) targeting the functional kinase activity of LRRK2 is a promising therapeutic approach (3). At least some pathogenic mutations, including the common G2019S mutation, enhance LRRK2 kinase activity and blocking kinase activity pharmacologically or genetically prevents LRRK2 neurotoxicity *in vitro* and in animal models (4-21). While aberrant LRRK2 kinase activity has been shown to affect a number of processes including cytoskeletal regulation (22-27), vesicular trafficking (15, 28, 29), autophagy (30-42), mitochondrial function (38, 43-45) and protein synthesis (9, 14, 46), precisely how LRRK2 mutations cause neuronal death remains unclear. Ultimately, prevention of LRRK2 pathogenesis will likely require a detailed understanding of the key mechanisms driving neurodegeneration. The clinical and pathological overlap between LRRK2-linked PD and idiopathic PD supports the notion that this understanding will have broad impact on our insight into PD even beyond cases where LRRK2 mutations are involved.

Penetrance of the LRRK2 G2019S mutation is incomplete even at advanced age, with estimates ranging from 25-80% and possibly varying among different ethnic populations (47-52). While some of this incomplete penetrance may be due to environmental factors, it is likely that background genetic variation impacts the probability of a LRRK2 G2019S carrier developing disease. Genetic variation can be harnessed as a powerful tool to identify which genes modify disease outcomes (53-57). This can be achieved effectively and efficiently using animal disease models when appropriate genetic tools and disease-relevant traits are available for study (58-61). In *Drosophila* for example, flies expressing human LRRK2 G2019S exhibit an age-dependent loss of dopamine neurons leading to pronounced locomotor deficits (14, 62). Importantly, flies expressing human wild-type LRRK2 do not exhibit similar PD-related phenotypes, supporting the existence of mutation-specific effects (14, 62). To determine the effects of genetic variation on LRRK2 G2019S-associated locomotor deficits, we used the *Drosophila* Genetic Reference Panel (DGRP), a large set of genetically-diverse background strains with fully-sequenced genomes (60). This panel has successfully been used to identify genetic modifiers in a number of diseases, infection, and cell stress models (63-70). We hypothesized that by assessing the effect of natural genetic variation on the LRRK2 G2019S locomotor phenotype which correlates with neurodegeneration in *Drosophila*, we could identify new candidate modifier genes that would provide important insights into the principle mechanisms underlying neuronal death.

## Materials and Methods

### Drosophila stocks and culture

There were 148 DGRP lines used in the study (Supplementary Material, Table S3) that were obtained from the Bloomington *Drosophila* Stock Center. The *UAS-LRRK2-G2019S* and *UAS-LRRK2* lines have been characterized elsewhere (14, 62) and were a gift from W. Smith. The double transgenic *Ddc-GAL4; UAS-LRRK2-G2019S* and *Ddc-GAL4; UAS-LRRK2* lines were created using *Ddc-GAL4* from the Bloomington *Drosophila* Stock Center (line 7010). Lines carrying RNAi constructs under UAS enhancer control were from the Bloomington *Drosophila* stock center (line numbers in parenthesis): *mAChR-C* (61306), *CG6420* (36088), *CG9003* (31362), *Eip63E* (34075), *IP3K2* (55240), *Khc-73* (38191), *nonC* (31094), *zfh1* (43195), *chi* (31049), *ct* (29625), *kon* (31584), *kay* (31322), *kn* (31916), *pbl* (28343), *pros* (26745) *smal* (55907) and control (36303) or Vienna *Drosophila* RNAi Center (line number in parenthesis): *CG12224* (31700), *CG14355* (109776), *CG14881* (110294), *CG17565* (110646), *CG33506* (107235), *Tie* (26879), *wry* (8021), *cv-c* (100247), *tup* (45859) and control (60100). All flies were reared and aged at 25°C/60% relative humidity under a 12 h light-dark cycle on standard cornmeal medium.

### Negative geotaxis

#### DGRP screening

Cohorts of 75-100 female flies (0-3 days-old, selecting against flies with visible signs of recent eclosion) collected from crosses between *Ddc-GAL4; UAS-LRRK2-G2019S* or *Ddc-GAL4; UAS-LRRK2* and each of 148 DGRP genetic backgrounds or *w*^*1118*^ were collected under brief anesthesia and transferred to fresh food vials to recover (25 flies/vial). Female *Ddc-GAL4; UAS-LRRK2-G2019S* expressing flies were used as they exhibit more pronounced locomotor deficits than males. We reasoned that for candidate suppressors, this would allow for a finer assessment of degree of suppression. Flies were aged for 6 weeks with transfer to fresh food twice per week. After 2 weeks, flies were transferred to empty vials, allowed 1 min to rest and then tapped to the bottom of the vial 3 times within a 1 second interval to initiate climbing. The position of each fly was captured in a digital image 4 s after climbing initiation using a fixed camera. Flies were then returned to their original food vial. Flies were subsequently tested again with the same protocol after 6 total weeks of aging. Automated image analysis was performed on digital images using the particle analysis tool on Scion Image to derive x-y co-ordinates for each fly thus providing the height climbed, as previously described (89). The performance of flies in a single vial was calculated from the average height climbed by all flies in that vial to generate a single datum (*N* =1). Performance of each line was then derived from the average scores of 3-4 vials tested for the line (*N* = 3-4).

#### RNAi screening

For testing effects of candidate modifiers on LRRK2 G2019S phenotypes, *Ddc-GAL4; UAS-LRRK2-G2019S* flies were crossed to each RNAi line. Additionally, *Ddc-GAL4; UAS-LRRK2-G2019S* or *UAS-LRRK2-G2019S* flies were crossed to an RNAi background line. For testing the effects of RNAi in the absence of LRRK2 G2019S, *Ddc-GAL4* flies were crossed to each RNAi line or to an RNAi background line. Cohorts of 75-100 female flies (0-3 days-old selecting against flies with visible signs of recent eclosion) were collected under brief anesthesia and transferred to fresh food vials to recover (25 flies/vial). Flies were aged for 2 and 6 weeks and negative geotaxis performance was measured as described above. Performance of TRiP and VDRC controls crossed to *Ddc-GAL4; UAS-LRRK2-G2019S* were almost identical and as the majority of RNAi stocks are from the TRiP collection, that RNAi control is shown.

### Dopamine neuron viability

*Ddc-GAL4; UAS-LRRK2-G2019S* flies were crossed to each RNAi line. *Ddc-GAL4; UAS-LRRK2-G2019S* or *UAS-LRRK2-G2019S* flies were crossed to an RNAi background line. Cohorts of 50 male and female flies (0-3 days-old) were collected under brief anesthesia and transferred to fresh food vials to recover (25 flies/vial). Flies were aged for 6 weeks then brains were harvested, fixed and permeabilized. Immunohistochemistry for tyrosine hydroxylase (TH) expressing neurons was performed using methods described elsewhere (90). Briefly, after blocking the brains in 5% normal goat serum/0.3% PBS-T, brains were incubated in anti-TH antibody (Immunostar) for 2 nights at 4 °C on a nutator. Brains were washed extensively, then incubated in alexa-fluor 488 anti-mouse secondary antibody for 2 nights at 4 °C on a nutator. Brains were washed extensively, mounted and imaged on a Zeiss LSM710 confocal microscope. Confocal z-stacks were acquired at 1 μm slice intervals. In order to quantify dopamine neurons, projection images through the brain to capture PAL (protocerebral anterior lateral), PPM1/2 (protocerebral posterior medial 1/2), PPM3, PPL1 (protocerebral posterior lateral 1) and PPL2 clusters were generated.

### Cytoscape functional network analysis of candidate modifier genes

Gene enrichment was assessed using the Cytoscape plug-in ClueGO, searching for GO molecular function and biological process terms. P values for group and term enrichment are calculated by Fisher Exact Test and corrected by Bonferroni step-down.

### Genome-wide association

Association analysis was performed as previously described (64). DGRP genotypes were downloaded from the website, http://dgrp.gnets.ncsu.edu/. Variants were filtered for minor allele frequency (≥ 0.05), and non-biallelic sites were removed. A total of 1,866,414 SNPs were included in the analysis. Negative geotaxis score for 148 DGRP/LRRK2 G2019S or 142 DGRP/LRRK2 (wildtype) F1 progeny were regressed on each SNP. To account for cryptic relatedness (91), GEMMA (v. 0.94) (92) was used to both estimate a centered genetic relatedness matrix and perform association tests using the following linear mixed model (LMM):

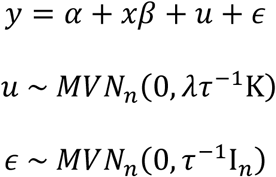

where, as described and adapted from Zhou and Stephens, 2012 (92), *Y* is the *n*-vector of negative geotaxis scores for the *n* lines, *α* is the intercept, *x* is the *n*-vector of marker genotypes, *β* is the effect size of the marker. *u* is a *n* x *n* matrix of random effects with a multivariate normal distribution (*MVN*_*n*_) that depends on *λ*, the ratio between the two variance components, *τ*^*–1*^, the variance of residuals errors, and where the covariance matrix is informed by K, the calculated *n* x *n* marker-based relatedness matrix. K accounts for all pairwise non-random sharing of genetic material among lines. *ϵ*, is a *n*-vector of residual errors, with a multivariate normal distribution that depends on *τ*^*–1*^ and I_*n*_, the identity matrix. Quantile-quantile (Q-Q) plot analysis indicated an appropriate fit to the LMM (Supplementary Materials, Fig. S3). Genes were identified from SNP coordinates using the BDGP R54/dm3 genome build. A SNP was assigned to a gene if it was within the gene’s transcription boundaries or no more than 1,000 bp upstream/downstream of those boundaries.

### Statistical analysis

Unless otherwise described in the methods, data were analyzed by ANOVA with Bonferroni correction for multiple comparisons.

### Data availability

SNP location data in the DGRP collection (Freeze 2.0 calls) are archived on the GSA Figshare portal and can additionally be downloaded from the DGRP data portal (http://dgrp2.gnets.ncsu.edu/data/website/dgrp2.tgeno).

## Results

### Locomotor dysfunction in LRRK2 G2019S flies is affected by genetic background

Flies expressing pathogenic LRRK2 G2019S via the dopaminergic *Ddc-GAL4* driver exhibit an age-dependent loss of dopamine neurons, with accompanying deficits in locomotor function (14, 62). We crossed the LRRK2 G2019S mutation into the suite of DGRP fly lines in order to examine the effect of natural genetic background variation on age-related locomotor dysfunction induced by mutant LRRK2. Double transgenic flies (*Ddc-GAL4; UAS-LRRK2-G2019S*) were crossed to 148 DGRP lines to produce F1 progeny deriving half of their genetic background from the parental DGRP line, permitting the assessment of dominant background effects on the LRRK2 G2019S locomotor phenotype.

In a w^1118^ background, LRRK2 G2019S locomotor deficits are absent in young flies and manifest around 6 weeks of age (14). We aged F1 progeny for 6 weeks and measured performance in negative geotaxis longitudinally at 2- and 6-weeks of age (Fig. 1). DGRP background has a strong effect on locomotor performance at both ages, (2 weeks, p <1.08 × 10^−66^; 6 weeks, p <4.04 × 10^−44^) and there is a significant interaction between age and DGRP background effects on geotaxis performance (p<0.0001), suggesting that the DGRP genetic backgrounds result in a distinct effect on locomotor performance in aged animals where G2019S locomotor deficits have previously been observed (14, 62). As the DGRP genetic backgrounds have not previously been tested for age-related locomotor function to our knowledge, we considered the possibility that variability in DGRP line locomotor performance could be due to the effects of aging on the different DGRP backgrounds and thus could occur independently of the LRRK2 G2019S mutation. To directly assess this, we carried out a control experiment in which the DGRP genetic background lines were crossed to a wildtype LRRK2-expressing line (*Ddc-GAL4; UAS-LRRK2*). F1 progeny from these crosses were aged for 6 weeks and tested for locomotor performance as the DGRP/LRRK2 G2019S lines were. The results reveal poor correlation between DGRP line means from the wildtype LRRK2 F1 progeny and the LRRK2 G2019S F1 progeny (Supplementary Material, Fig S1). Additionally, there were no common GWAS candidate genes nominated by the wildtype LRRK2 and LRRK2 G2019S data sets when considering SNPs associated at a p ≤ 10^−6^threshold (Table 2). Hence, the effect of genetic background on locomotor performance does not occur independently of LRRK2 G2019S.

**Figure 1:**
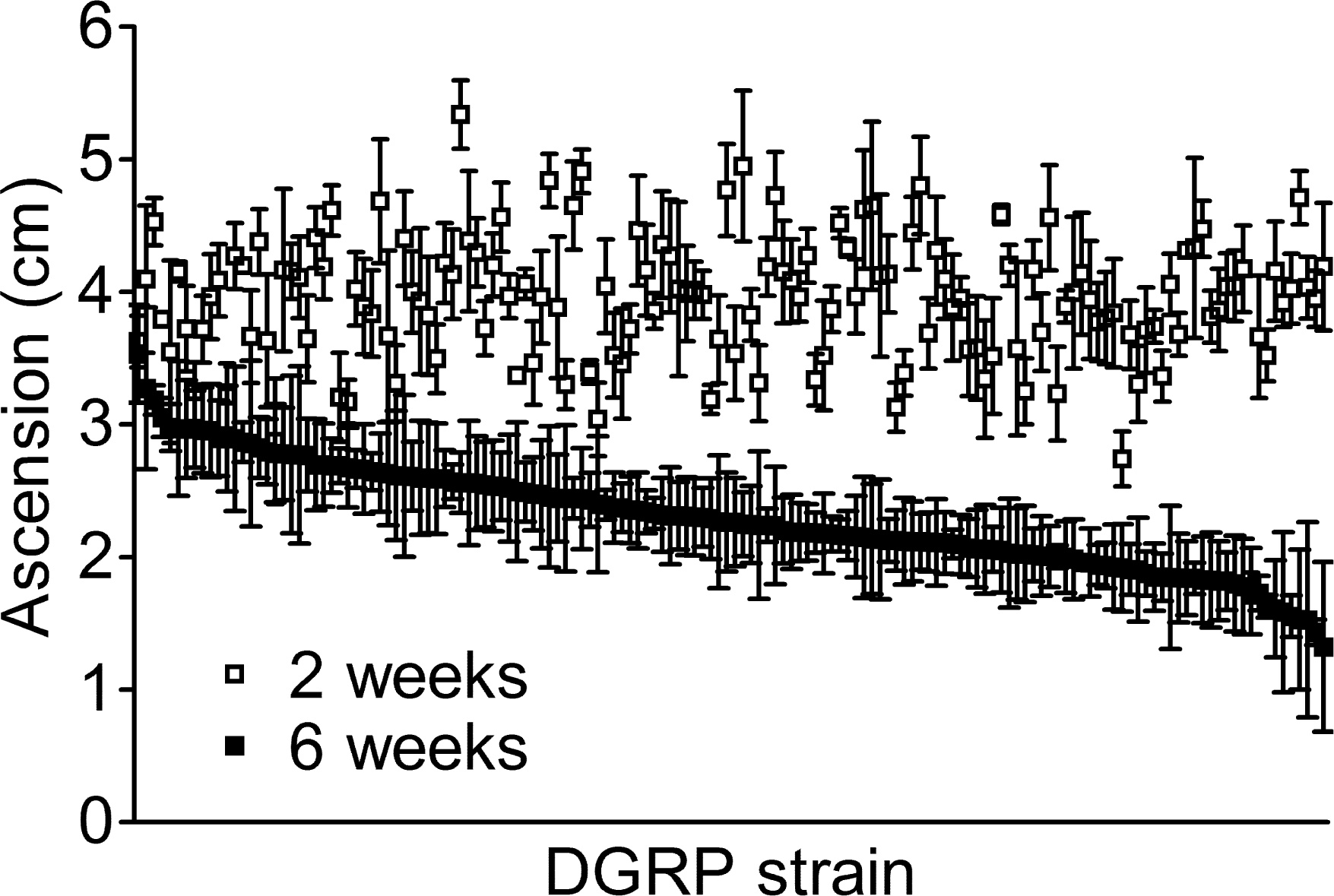
Locomotor function of LRRK2 G2019S flies depends on genetic background. Negative geotaxis of *Ddc-GAL4; UAS-LRRK2-G2019S* flies in 148 DGRP backgrounds at 2 and 6 weeks of age. DGRP background has a strong effect on locomotor performance at both ages (2 weeks, p <1.08 × 10^−66^; 6 weeks, p <4.04 × 10^−44^).

### Genome-wide association reveals candidate modifiers of LRRK2 G2019S locomotor dysfunction

The large range of geotaxis scores seen in LRRK2 G2019S flies with different DGRP backgrounds suggests that genetic variation influences mutant LRRK2-mediated locomotor dysfunction. Quantitative geotaxis scores of aged flies (6-week-old) were used to test for association with genome-wide single nucleotide polymorphisms (SNPs) in an unbiased manner. We tested whether a given SNP was associated with the LRRK2 G2019S locomotor phenotype using a genome-wide association approach. With a large number of SNPs (1,866,414) tested over just 148 lines, the study was not sufficiently powered for SNP associations to survive multiple testing correction. However, our goal was not to treat these associations as being definitive, but to use these SNPs to nominate candidate modifier genes for subsequent functional validation. This approach yields a number of genes that potentially underlie the variable expressivity of LRRK2 G2019S phenotypes for further testing.

In total, 177 candidate genes collectively harboring 280 SNPs are associated with variability in the LRRK2 G2019S locomotor phenotype at a p-value threshold of p < 10^−5^ (Supplementary Material, Table S1). A subset of 19 genes (with 31 associated SNPs) meet the more restrictive nominal p-value threshold of p < 10^−6^, and can therefore be considered top candidates by statistical association (Table 1). These genes are hereafter referred to as top association candidates. Within this group, 15 genes have human orthologs (Table 1), and several genes harbor multiple associated SNPs (4 in *wry* and 2 each in *CG43277* and *tie*). These genes cover a broad range of annotated functions, including microtubule binding and motor activity for *Khc-*73 (71), motor neuron axon guidance for *zfh1* (72, 73), notch signaling for *wry* (74) and calcium signaling for *IP3K2* (75). *KIF13A*, one of the putative human orthologs of *Khc-73*, is reported to play a role in the trafficking of mannose-6-phosphate-containing vesicles from the trans-Golgi to the plasma membrane (76). This is notable given the proposed role for LRRK2 in regulating vesicular trafficking through its interaction with Rabs (28, 29). A gene ontology enrichment analysis on all 177 candidates revealed an enrichment of genes within a number of functional groups, the most significant being dendrite guidance (p = 2.9 × 10^−6^) and regulation of cell projection organization (p = 3.0 × 10^−5^) (Supplementary Material, Table S2). These findings are striking considering the recognized role of LRRK2 in regulating neurite outgrowth as well as the loss of neurite complexity consistently reported in cultured neurons or animals expressing LRRK2 G2019S (22-27, 77-81). Collectively, we termed genes from the dendrite guidance and regulation of cell projection GO categories as neuron projection candidates (Table 3). None of these candidates fall within the top association genes. Performing gene enrichment analysis exclusively on the list of top association genes does not give any functional enrichment, which is not surprising given that there are only 19 genes.

**Table 1:** Top association candidate genes

| Rank | SNP<sup>a</sup> | Candidate gene(s) | p-value | FDR | Human ortholog | OMIM |
| --- | --- | --- | --- | --- | --- | --- |
| 1 | 3R:9973680 | *CG14355* | 2.65E<sup>-07</sup> | 0.37 | – | – |
| 2 | 2R:10746081 | *CG33506* | 3.99E<sup>-07</sup> | 0.37 | *TMEM177* | – |
| 3 | 2R:15525692 | *CG43277<sup>b</sup>* | 2.46E<sup>-06</sup> | 0.47 | – | – |
| 4 | 2R:11417294 | *Khc-73* | 2.87E<sup>-06</sup> | 0.47 | *KIF13A/KIF13B* | 605433/607350 |
| 5 | X:14035579 | *CR44658<sup>b</sup>* | 3.17E<sup>-06</sup> | 0.47 | – | – |
| 6 | 3R:26611553 | *zfh1* | 3.69E<sup>-06</sup> | 0.47 | *ZEB2* | 605802 |
| 7 | 2L:1844384 | *wry* | 4.10E<sup>-06</sup> | 0.47 | *DNER* | 607299 |
| 8 | 3L:4510450 | *Tie* | 4.12E<sup>-06</sup> | 0.47 | *TEK/Tie1* | 600221/600222 |
| 9 | X:13199667 | *IP3K2* | 4.41E<sup>-06</sup> | 0.47 | *ITPKA/ITPKB* | 147521 |
| 10 | 3R:22549408 | *CG6420* | 5.15E<sup>-06</sup> | 0.47 | *WDR20* | – |
| 11 | 3R:7806766 | *CG12224* | 5.38E<sup>-06</sup> | 0.47 | *KCNAB2* | 601142 |
| 12 | 3R:12260571 | *CG17565/CG14881* | 6.16E<sup>-06</sup> | 0.47 | *FNTB/ARMC7* | 134636 |
| 13 | 3R:13196662 | *CR44256<sup>b</sup>* | 6.31E<sup>-06</sup> | 0.47 | – | – |
| 14 | 3L:3521221 | *Eip63E* | 7.42E<sup>-06</sup> | 0.47 | *CDK14* | 610679 |
| 15 | X:6727165 | *nonC/mAChR-C* | 7.58E<sup>-06</sup> | 0.47 | *SMG1/GPR119* | 607032/300513 |
| 16 | 2R:7528497 | *CG9003* | 7.61E<sup>-06</sup> | 0.47 | *FBXL20* | 609086 |
| 17 | 3R:18824405 | *Nha2<sup>b</sup>* | 8.20E<sup>-06</sup> | 0.49 | *SLC9B1/SLC9B2* | 611527 |

<sup>a</sup>SNP with most statistically significant association for the candidate gene
<sup>b</sup>No RNAi line available for testing candidate gene

**Table 2:**
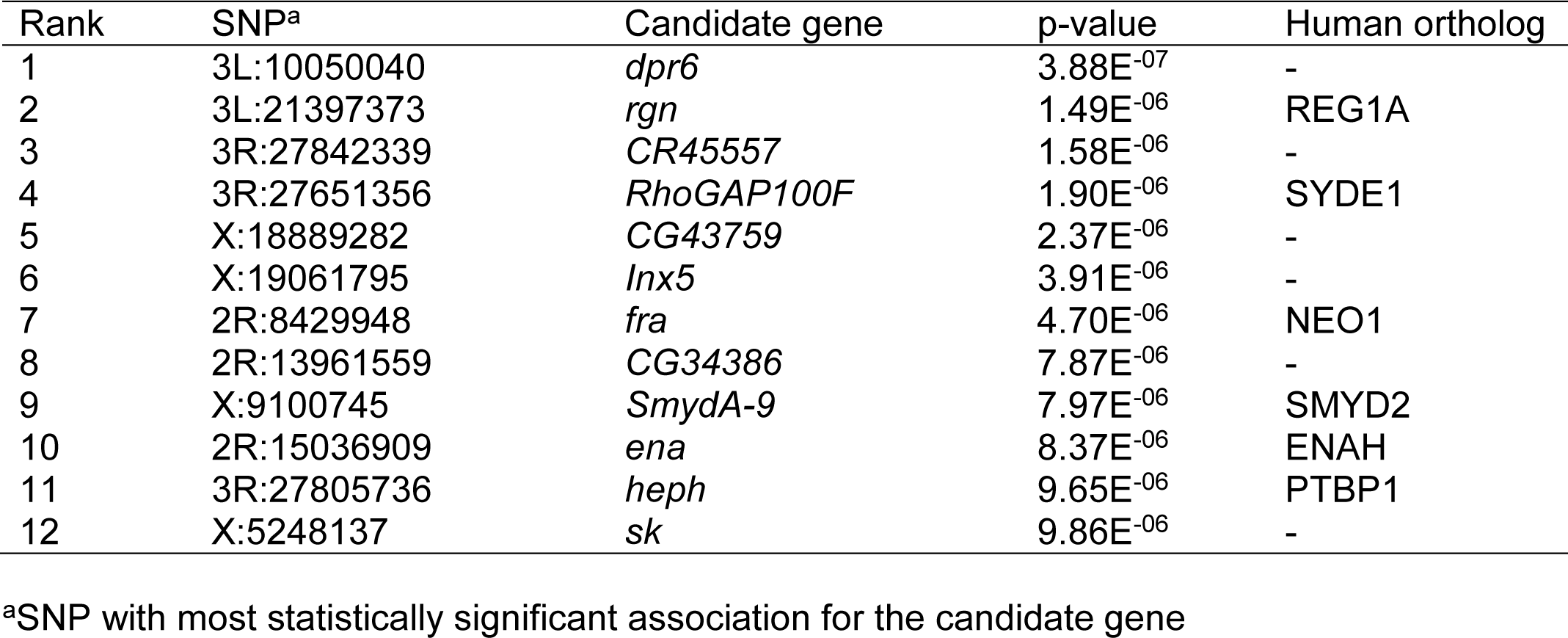
Wild type LRRK2 top association candidate genes

**Table 3:** Neuron projection candidate genes

| Candidate gene | SNP<sup>a</sup> | p-value | Human ortholog | OMIM |
| --- | --- | --- | --- | --- |
| *chi* | 2R:19919006 | 1.67E<sup>-05</sup> | — | — |
| *ct* | X:7535575 | 6.06E<sup>-05</sup> | *CUX1/CUX2* | 116896/610648 |
| *cv-c* | 3R:10260649 | 7.42E<sup>-05</sup> | *DLC1* | 604258 |
| *kay* | 3R:25611636 | 7.16E<sup>-05</sup> | *FOS/FOSL1/FOSL2* | 164810/136515/601575 |
| *kn* | 2R:10671765 | 9.12E<sup>-05</sup> | *EBF3* | 607407 |
| *kon* | 2L:18491720 | 6.20E<sup>-05</sup> | *CSPG4* | 601172 |
| *pbl* | 3L:7905741 | 1.48E<sup>-05</sup> | *ECT2* | 600586 |
| *pros* | 3R:7155471 | 6.68E<sup>-05</sup> | *PROX1* | 601546 |
| *smal* | 2L:6187849 | 4.05E<sup>-05</sup> | *DDR1/DDR2* | 600408/191311 |
| *tup* | 2L:18863703 | 1.83E<sup>-05</sup> | *ISL1* | 600366 |

<sup>a</sup>SNP with most statistically significant association for the candidate gene

### Targeted knock-down of candidate genes impacts LRRK2 G2019S neurodegeneration

In order to test the candidate modifier genes nominated by our initial screen, we pursued the top association candidates and neuron projection candidates (25 genes total) to test their ability to influence the LRRK2 G2019S locomotor phenotype. Inclusion of the neuron projection candidates was on the basis of compelling functional evidence linking LRRK2 G2019S to altered neurite morphology (22-27, 77-80) as well as our hypothesis that we may identify novel modifiers of LRRK2 G2019S-mediated neurodegeneration that could provide insight into pathogenesis. We used RNAi to probe whether lowering candidate gene expression influenced the LRRK2 G2019S locomotor phenotype. RNAi lines for each candidate were mated to *Ddc-GAL4; UAS-LRRK2-G2019S* flies to generate progeny with concomitant LRRK2 G2019S expression and candidate modifier knock-down (KD) in dopamine neurons. RNAi lines were available for all candidates except two non-protein-coding genes (*CR44658* and *CR44256*), *CG43277* and *nha-2*. Although this simple loss-of-function approach does not fully assess the contribution of each gene to the phenotype, it allows us to directly probe for a phenotype modifying effect of candidate gene KD specifically within dopamine neurons while using readily available genetic tools.

As with the initial genetic screen, we assessed the LRRK2 G2019S locomotor phenotype by measuring negative geotaxis in 2- and 6-week old flies. RNAi for *CG9003, cv-c* and *kn* each result in reduced geotaxis performance in 2 week-old flies, raising the possibility that they may impact locomotor function independently of LRRK2 G2019S (Fig. 2). Indeed, *Ddc-GAL4* driven KD of these three candidate genes without LRRK2 G2019S expression induce locomotor dysfunction in aged flies (Supplementary Material, Fig S2). Knocking down 7 of the 15 top association candidates tested (47%) significantly enhances LRRK2 G2019S locomotor deficits at 6 weeks of age (*mAChR-C, CG14355, CG14881, CG17565, CG33506, CG9003,* and *tie*). None of the 15 top association candidates tested suppressed the G2019S LRRK2 locomotor phenotype when knocked down. RNAi for 6 of the 10 neuron projection candidates (60%) significantly affects the LRRK2 G2019S phenotype at 6 weeks of age (Fig. 2). Of these 6, 5 candidates (*cv-c, kn, pbl, pros* and *tup*) enhance and 1 candidate (*ct*) suppresses the phenotype when knocked-down.

**Figure 2:**
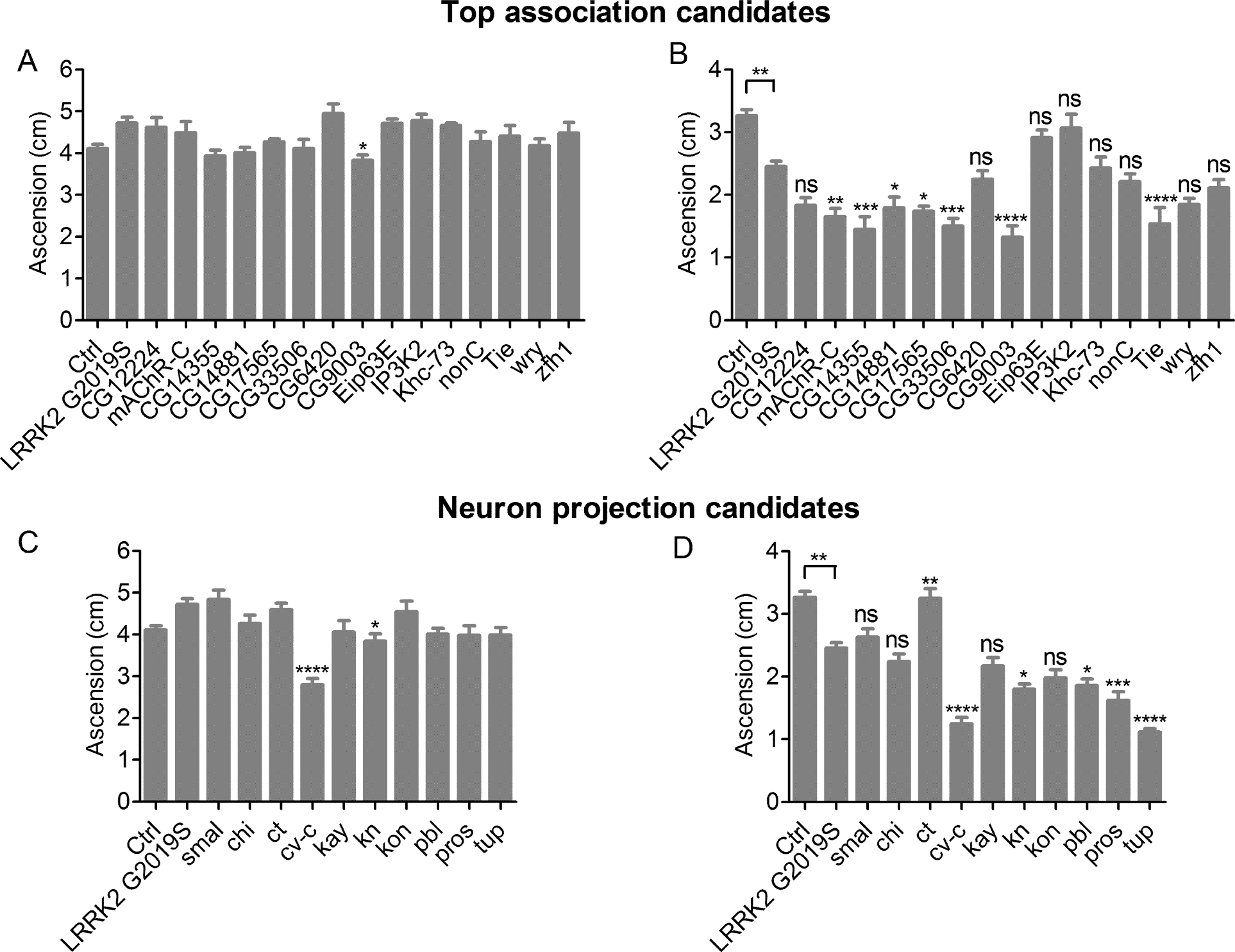
Effect of candidate modifier gene knock-down on LRRK2 G2019S locomotor dysfunction. Top association candidates were tested at 2 weeks (A) and 6 weeks (B) of age. Neuron projection candidates were tested at 2 weeks (C) and 6 weeks (D) of age. In each case, there was a significant effect of genotype on geotaxis (individual ANOVAs, p < 0.0006). In 6-week-old flies, Bonferroni post-tests revealed a significant effect of G2019S LRRK2 expression and an enhancement or suppression effect of RNAi for some candidate genes on the performance of G2019S LRRK2-expressing flies (ns not significant, * p <0.05, ** p < 0.01, *** p <0.001, **** p <0.0001). Ctrl is *+/+; UAS-LRRK2-G2019S/+* and LRRK2 G2019S is *Ddc-GAL4/+; UAS-LRRK2-G2019S/+*.

To test whether a significant candidate modifier KD effect on the LRRK2 G2019S locomotor phenotype correlates with a change in neuronal degeneration, we next examined dopamine neuron viability in aged flies for the RNAi lines which modified locomotor performance in aged LRRK2 G2019S flies, but which had no effect in the absence of LRRK2 G2019S. Hence, *CG9003, cv-c* and *kn* were not advanced to this stage of testing. We assessed five dopamine neuron clusters (PAL, PPM1/2, PPM3, PPL1 and PPL2) in the fly brain that are readily quantifiable. Most of the top association candidates have a modest effect on dopamine neuron viability; the only candidate reaching statistical significance is *CG33506*, which is the second highest ranked candidate based on SNP association. While LRRK2 G2019S expression alone causes a 20% loss of dopamine neurons for the five clusters counted, additional *CG33506* KD leads to a 43% decrease. This correlates well with locomotor effects, where geotaxis scores decrease by 25% with LRRK2 G2019S expression alone and 54% with additional *CG33506* KD. *CG33506* is uncharacterized, although reported to be expressed in fly heads (82). The human ortholog of *CG33506* is *TMEM177*, which has 42.1% identity at the nucleotide level and 25.3% identity at the protein level. TMEM177 is a multi-pass transmembrane protein, largely uncharacterized but expressed throughout the body, including in brain, and thought to localize to mitochondria (83).

From our neuron projection candidates subject to dopamine neuron assessment, 3 out of 4 of the RNAi lines exhibit a significant modifier effect on neuronal viability (Fig. 3). In accordance with their effects on the LRRK2 G2019S locomotor phenotype, both *pros* and *pbl* enhance loss of dopamine neurons upon KD, while *ct* suppresses it. KD of *tup* also enhances LRRK2 G2019S-mediated dopamine neuron death consistent with its effects on geotaxis performance, although the difference did not reach statistical significance. All 3 neuron projection candidates that significantly modify both locomotor and dopamine neuron phenotypes when knocked-down have human homologs and established roles in neurite morphogenesis. While RNAi effects on locomotor function and dopamine neuron viability in LRRK2 G2019S flies generally correlate, it should be noted that KD of 3 genes (*mAChR-C, tie* and *CG14355*; 30% of candidates tested) produce non-significant effects on dopamine neuron viability that do not correspond directionally with their effects on geotaxis (Fig. 3).

**Figure 3:**
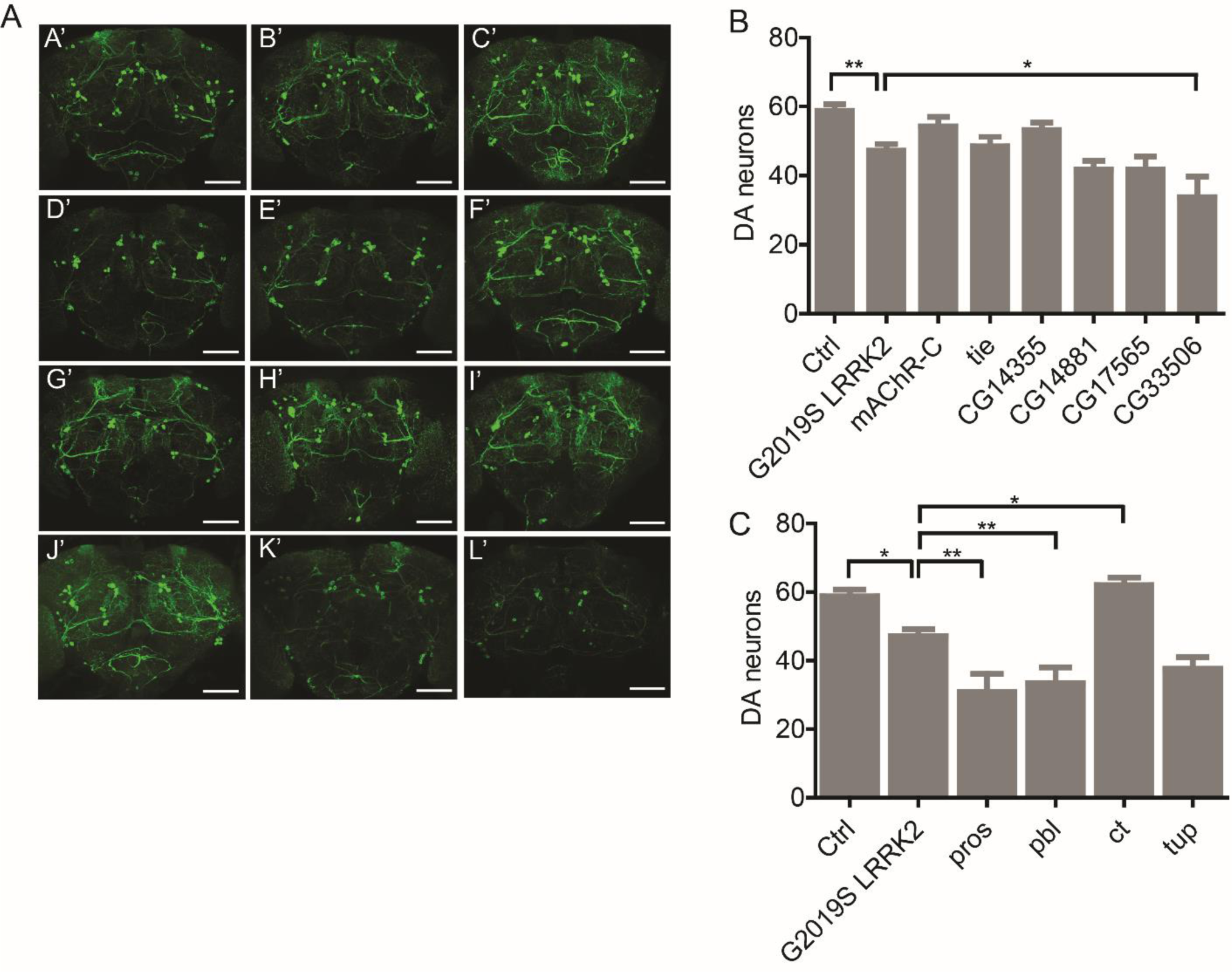
Effect of candidate modifier gene knock-down on LRRK2 G2019S dopamine neuron loss. (A) Confocal projection images through the brain of control +/+; UAS-LRRK2-G2019S/+ (A’), and *Ddc-GAL4/+; UAS-LRRK2-G2019S/+* (B’) flies and G2019S LRRK2 expressing flies with RNAi-mediated knockdown of mAChR-C (C’), tie (D’), CG14355 (E’), CG14881 (F’), CG17565 (G’), CG33506 (H’), pros (I’), pbl (J’), ct (K’) and tup (L’). Scale bars are 60 μM. Quantitation of total dopamine neurons in five clusters (PPM1/2, PPM3, PPL1, PPL2 and PAL) for top association (B) and neuron projection (C) candidate genes. There was a significant effect of genotype for both (individual ANOVAs, p < 0.0001) and Bonferroni post-tests revealed a significant effect of G2019S LRRK2 expression and knock-down of *CG33506, pros, pbl* and *ct* candidate modifiers (* p <0.05, ** p <0.01).

## Discussion

Genetic background can be an important factor in determining whether disease manifests in an individual harboring a given pathogenic mutation. LRRK2 G2019S penetrance is age-related but incomplete even by the eighth decade (47-52, 84). Environmental and genetic factors are both likely to play a role in determining whether a LRRK2 G2019S carrier develops PD. In *Drosophila*, the impact of natural genetic variation on a given trait can be assessed in an unbiased manner using established genetic tools such as the DGRP suite of flies. The DGRP facilitates assessment of quantifiable phenotypes in a large number of fully-sequenced backgrounds, and has been used on numerous occasions to demonstrate variation in disease-relevant traits (60, 64-70). We used the DGRP to show that genetic background strongly influences the age-related locomotor phenotype seen in LRRK2 G2019S expressing flies, and identified candidate modifier genes of mutant LRRK2. Our unbiased genome-wide association study identified numerous SNPs associated with the LRRK2 G2019S locomotor phenotype. Importantly, the genes nominated may affect the LRRK2 G2019S locomotor phenotype not just as downstream mediators of LRRK2 neurotoxicity, but also as upstream regulators of LRRK2 expression/function or by acting in parallel pathways that functionally converge with LRRK2 to influence dopamine neuron integrity. As anticipated, there was no strong correlation between the DGRP line effects at 2 weeks and at 6 weeks of age in our study (Fig. 1). Genetic background can have distinct influences on locomotor performance in young and aged flies, manifest as a relatively high-performing background at 2 weeks that becomes a relatively low-performing background at 6 weeks, or vice versa (85). This is likely due to distinct, and in some cases even opposing effects of genetic background on performance vigor in young flies and on age-related changes that occur over this large portion of the fly life span. Hence, the complex interplay between genetic background and aging does not produce strongly correlative performance at 2 and 6 weeks of age.

Our GWAS relies on the association of many SNPs over just 148 lines, which for the locomotor phenotype analysis performed did not enable sufficient power to identify definitive SNPs with very high association that survive multiple testing correction. Since this increases the possibility of identifying false-positives, we used the SNPs simply to nominate candidate modifier genes for subsequent detailed validation. Based on this simple goal, we did not examine the molecular nature of the associated SNPs in terms of how they might impact expression or function of candidate genes. The vast majority of SNPs in these candidate genes were non-coding and therefore it is difficult to predict how they may impact expression levels. Further, obtaining definitive evidence to address this question would require experimental testing of the SNPs and gene function, which was beyond the goals of the study.

Considering the possibility that phenotypic variability in the DGRP backgrounds could occur independently of LRRK2 G2019S, and alternatively be related to effects of the different backgrounds or LRRK2 overexpression on age-related negative geotaxis behavior, we tested a separate cohort of F1 progeny using the same genetic backgrounds but this time expressing wild type LRRK2. This revealed poor correlation between the DGRP backgrounds expressing LRRK2 with those expressing LRRK2 G2019S (Supplementary Material, Fig S1). We further performed a GWAS on the wild-type LRRK2 cohort, deriving a total of 12 top association candidates (at p ≤10^−6^), none of which overlapped with the top association candidates obtained via a LRRK2 G2019S GWAS (Tables 1 and 2). These lines of evidence argue that the phenotypic variability observed via LRRK2 G2019S is not simply due to effects of the DGRP backgrounds or LRRK2 expression on age-related locomotor performance and is dependent on mutant LRRK2 G2019S expression. Attempting to undertake a joint analysis in which the deviation in performance between LRRK2 and LRRK2 G2019S for each genetic background forms a basis for GWAS analysis was precluded by the testing of each F1 progeny cohort separately. A complex behavioral phenotype such as negative geotaxis is prone to cohort effects caused by environmental variables that occur over the course of aging. These additional variables confound the ability to meaningfully isolate “mutation-specific” effects by subtracting performance scores between two cohorts that are reared, aged and tested separately. These possible cohort effects could in theory also impact the correlation analysis of DGRP backgrounds expressing LRRK2 vs.

LRRK2 G2019S, yet likely not in a manner that would prevent our ability to observe a strong correlation between the 2 sets. Additionally, while we were not able to incorporate an integrated analysis of this nature, the approach we took by performing a GWAS directly on the LRRK2 G2019S cohort is supported by the subsequent testing and observation that 13 out of 25 top association and neuron projection candidates nominated by this GWAS were validated as modifiers of locomotor function in aged LRRK2 G2019S flies. Hence, we are confident that our approach has led to the identification of authentic LRRK2 G2019S phenotype modifiers that warrant further investigation.

Among the list of genes nominated by genome-wide association, we prioritized for further testing a subset of candidates with the strongest statistical association, and those in two enriched GO categories (dendrite guidance and regulation of cell projection organization) where direct relevance to LRRK2 function is supported by prior studies. By considering not only our top association hits but also gene ontology-enriched candidates associated at p < 10^−5^, we hoped to partially overcome the lack of statistical power for identifying definitive candidates based on SNP associations. This approach led to a pool of potential targets for which subsequent validation permitted identification of LRRK2 G2019S genetic modifiers of high confidence. Knocking down 13 out of these 25 candidates significantly modifies the LRRK2 G2019S locomotor phenotype in aged flies, and after subsequent testing of dopamine neuron viability, we derived a final set of candidate genes: *CG33506, pros, pbl* and *ct* that significantly modify both locomotor dysfunction and dopamine neuron loss phenotypes of LRRK2 G2019S in *Drosophila*. That we were ultimately able to identify a set of candidate modifier genes for both locomotor and dopamine neuron viability phenotypes supports the GWAS study in terms of the targets nominated by this approach. *pros* is a transcriptional regulator involved in neuronal differentiation and is required for axon and dendrite development. *pbl* is a Rho guanine nucleotide exchange factor (GEF) involved in a number of processes including axonogenesis. *ct* is a homeoprotein transcription factor known to influence dendritic morphology in the fly dendritic arborization neurons. A role for LRRK2 in regulating neurite outgrowth has been documented through the interaction with proteins important in cytoskeletal dynamics such as Rac1, PAK6, ERM proteins, MARK1 and Tau as well as the cytoskeletal proteins Actin and Tubulin themselves (22-27, 77-80). Studies have shown that these cytoskeletal protein interactions may be perturbed by the G2019S mutation leading to neuronal injury *in vitro* (22-27, 77-80). Our study reinforces this relationship and crucially goes further by demonstrating a direct link between genes important for neurite development/maintenance and DA neurodegenerative phenotypes *in vivo*. Further, we have identified novel candidate interactors for LRRK2 that, through further characterization, may help delineate the precise mechanism by which LRRK2 regulates neurite morphogenesis.

It is currently hard to hypothesize how *CG33506* (human ortholog *TMEM177*) may be involved in LRRK2 biology or neurodegeneration. TMEM177 is thought to localize to mitochondrial membranes where protein interaction mapping suggests an interaction with numerous mitochondrial ribosomal proteins (83). Our previous work shows that LRRK2 impacts bulk protein synthesis which contributes to LRRK2 G2019S neurodegeneration through unclear mechanisms (14), but whether it affects mitochondrial protein synthesis is unknown. Mutations in another multi-pass transmembrane protein TMEM230 were recently associated with familial Parkinson’s disease (87). TMEM230 did not co-localize with a mitochondria marker but is thought to localize to vesicles and impact synaptic vesicle trafficking (87). Considering this and the broad range of functions encompassed by transmembrane proteins, it is unclear whether TMEM177 and TMEM230 have any connection relevant to LRRK2.

Marcogliese et al. recently conducted a genetic screen to identify modifiers of LRRK2 I2020T eye degeneration in *Drosophila* (88). They initially screened through deficiency lines (containing deletions that cover almost the entire genome) at 25°C and 29°C in flies expressing I2020T LRRK2 via the eye-specific *GMR-GAL4* driver, followed by functional testing of candidate modifiers. Their set of final candidate modifiers validated to impact dopamine neuron viability in addition to eye degeneration does not overlap with our candidates that modify both age-related locomotor dysfunction and dopamine neuron loss. One simple explanation for this is that modifiers for different pathogenic variants of LRRK2 may be distinct, although both G2019S and I2020T mutations are thought to be neurotoxic at least in part through an elevation of LRRK2 kinase activity (20). One possible explanation for this is that as a starting point, they expressed LRRK2 I2020T in the eye and used an eye degeneration phenotype for their screen while we expressed in LRRK2 G2019S in dopamine neurons and used an age-related locomotor phenotype which is directly related to dopamine neuron loss. We believe our approach has the advantage that age-related locomotor dysfunction due to dopamine neuron loss is potentially more directly related to PD than eye degeneration phenotypes. The higher consistency of effects on locomotor and dopamine neuron phenotypes in our study would also suggest this. Of the 10 candidates selected for dopamine neuron viability assessment in our study, 6 candidates (60%) affected both phenotypes in the same direction, whereas 6 out of 16 candidates (38%) showed consistent effects in their I2020T LRRK2 study. As noted by the authors, the dichotomy in directional effects between eye degeneration and dopamine neuron loss encountered may have been caused by false positive results in their initial screen.

Our findings highlight the influence of natural genetic variation in the expressivity of disease-related phenotypes. The DGRP has been successfully utilized to uncover the impact of genetic background variation on traits linked to retinitis pigmentosa and resistance to infection (69), along with cellular responses to ER stress (65) and oxidative stress (70). We have now extended this to locomotor deficits in a model of Parkinson’s disease and additionally identified novel candidate modifier genes for a disease pathway relevant to LRRK2. Effective prevention of neuronal death resulting from LRRK2 mutations will require a detailed understanding of the key mechanisms driving neurodegeneration. This study generates new targets to investigate and may direct future studies on LRRK2 pathobiology and therapeutic intervention in PD.

## Acknowledgments

We thank the Advanced Light Microscopy Core for subsidized confocal microscope use.

